# Orexin/hypocretin immunoreactivity in the lateral hypothalamus is reduced in genetically obese but not in diet-induced obese mice

**DOI:** 10.1101/145508

**Authors:** J. Antonio González, Jochen H. M. Prehn

## Abstract

The mechanisms that link diet and body weight are not fully understood. A diet high in fat often leads to obesity, and this in part is the consequence of diet-induced injury to specific hypothalamic nuclei. It has been suggested that a diet high in fat leads to cell loss in the lateral hypothalamus, which contains specific populations of neurones that are essential for regulating energy homoeostasis; however, we do not know which cell types are affected by the diet. We studied the possibility that high-fat diet leads to a reduction in orexin/hypocretin (O/H) and/or melanin-concentrating hormone (MCH) immunoreactivity in the lateral hypothalamus. We quantified immuno-labelled O/H and MCH cells in brain sections of mice fed a diet high in fat for up to 12 weeks starting at 4 weeks of age and found that this diet did not modify the number of O/H- or MCH-immunoreactive neurones. By contrast, there were fewer O/H- (but not MCH-) immunoreactive cells in genetically-obese *db/db* mice compared to wild-type mice. Non-obese, heterozygous *db/+* mice also had fewer O/H-immunoreactive cells. Differences in the number of O/H-immunoreactive cells were only a function of the *db* genotype but not of diet or body weight. Our findings show that the lateral hypothalamus is affected differently in genetic and in diet-induced obesity, and support the idea that hypothalamic neurones involved in energy balance regulation are not equally sensitive to the effects of diet.

## 1 Introduction

A diet high in fat often leads to obesity and type 2 diabetes in both humans and animals (Astrup *et al.*, 2008) but the mechanisms that link diet and metabolic disease are not fully understood. One consequence of high-fat diet intake is damage to the hypothalamus, which is an essential component of the network that regulates appetite and metabolism in the body (Waterson & Horvath, 2015). For example, inflammation and leptin resistance in the arcuate nucleus (Münzberg *et al.*, 2004; Thaler *et al.*, 2012), and reactive astrogliosis in several hypothalamic nuclei (Buckman *et al.*, 2013) have been observed in response to high-fat diet. In addition, Moraes *et al.* (2009) observed loss of neurones in the arcuate and in the lateral hypothalamus of animals fed a diet high in fat (although others have failed to see this effect; see Lemus *et al.*, 2015; Namavar *et al.*, 2012). Thus, it is possible that obesity and metabolic disease are the result of fat-induced hypothalamic injury that may include inflammation and neuronal loss (reviewed in Jais & Brüning, 2017).

Interestingly, much of the damage that is seen in the arcuate following high-fat diet seems to preferentially affect neurones involved in appetite regulation, namely orexigenic neurones that express neuropeptide Y and anorexigenic neurones that express the peptide pro-opiomelanocortin (Moraes *et al.*, 2009; Horvath *et al.*, 2010). In the lateral hypothalamus two neuronal populations that are modulated by nutrients and hormones (e.g. leptin and insulin) and that in turn regulate appetite, peripheral glucose, and adiposity express the peptides melanin-concentrating hormone (MCH) or orexin/hypocretin (O/H) (Burdakov *et al.*, 2013). However, we do not know if MCH and O/H cells of the lateral hypothalamus are specifically affected by a high-fat diet intake.

To investigate if diet modifies the number of MCH or O/H neurones we counted MCH- and O/H-immunoreactive (ir) cells in the hypothalamus of mice that were fed a diet high in fat. We also quantified MCH- and O/H-ir cells in a genetic model of obesity, the *db/db* mouse, to assess if any potential changes in cell counts were linked to the obese phenotype itself or specifically to the high-fat contents in the diet. Compared to control mice, diet-induced obese mice had a similar number of O/H- and MCHir cells. By contrast, *db/db* mice had fewer O/H- (but not MCH-) ir cells.

## 2 Materials and methods

### 2.1 Animals

Animal experiments were performed in accordance with the European Communities (Amendment of Cruelty to Animals Act 1876) Regulations 2002 under license from the Department of Health and Children, Ireland, and were approved by the Research Ethics Committee of The Royal College of Surgeons in Ireland (REC570).

Two animal models of obesity were used for these experiments: genetically obese *db/db* mice and diet-induced obese mice. Diet-induced obesity was set up with wild-type male mice (C57BL/6JOlaHsd, 3 weeks-old, *n* = 24) from Harlan Laboratories (Blackthorn, UK), housed in 6 cages of 4 animals each with free access to water and standard chow (Teklad 2018, 18% cal from fat). After one week (i.e. at 4 weeks of age) food in 3 of these cages was replaced by a diet high in fat (Research Diets D12492, 60% cal from fat), thus creating 3 control (standard diet) and 3 experimental (high-fat diet, HFD) groups. Eight animals (*n* = 4 control and *n* = 4HFD) were taken at weeks 4, 8, or 12 after introducing the diet high in fat for the experiments described here.

Genetically-obese *db/db* mice develop severe obesity and diabetes (Hummel *et al.*, 1966) because of a mutation in the leptin receptor which renders leptin signalling ineffective (Chen *et al.*, 1996). Heterozygous (*db/+*) mice are not different physiologically or morphologically from wild-type mice (Hummel *et al.*, 1966), and thus were used as controls. Male *db/db* (BKS.Cg-+*Lepr*^*db*^/+*Lepr*^*db*^/OlaHsd; *n* = 8) and *db/+* (BKS.Cg-*Dock7*^*m*^ +/+ *Lepr*^*db*^/OlaHsd; *n* = 8) mice were obtained from Harlan Laboratories, UK, at 7 weeks of age and were housed with free access to water and standard diet (Teklad 2018, as above). They were taken at 8 (*n* = 4from each group) or 14 (*n* = 4 from each group) weeks of age for the experiments described below.

### 2.2 Immunohistochemistry

Animals were deeply anaesthetised with pentobarbital sodium (20% solution, 0.1 ml i.p.) and then perfused transcardially with 20 ml phosphate-buffered saline (PBS) followed by 20 ml of paraformaldehyde (4% in PBS, pH 7.4). The brain was immediately removed and placed in paraformaldehyde (as above) for about two hours, then kept in sucrose solution (30% in PBS) for 48–72h at 4°C until the brain precipitated to the bottom of the container. Afterwards, the brain was frozen and stored at −80°C until further processing.

Coronal brain sections 30-μm thick were cut rostrocaudally (Fig. 1A) with a cryostat (Leica CM1950; Leica Biosystems, Nussloch, Germany) and stored in cryoprotectant (%v/v, glycerol 30, ethylene glycol 30, phosphate buffer 10, water 30) at −20°C. Sections spanning the full extent of the posterior hypothalamus were collected. Then, with the aid of a low-magnification microscope, the section at the caudal end of the optic chiasm (at the point where the optic tract begins its dorso-lateral trajectory towards the thalamus; Fig. 1B) was selected as ‘section 1’, and five more sections were collected every 6th in rostrocaudal sequence, i.e. we collected 6 sections per brain at 180 μm intervals (Fig. 1A). (The seventh section in this same sequence would have had no O/H cells.) That way we ensured that the sections were histologically equivalent at each level across all animals, and that we sampled along the full extent of the O/H field in the hypothalamus (see e.g. Hahn, 2010).

**Figure 1.**
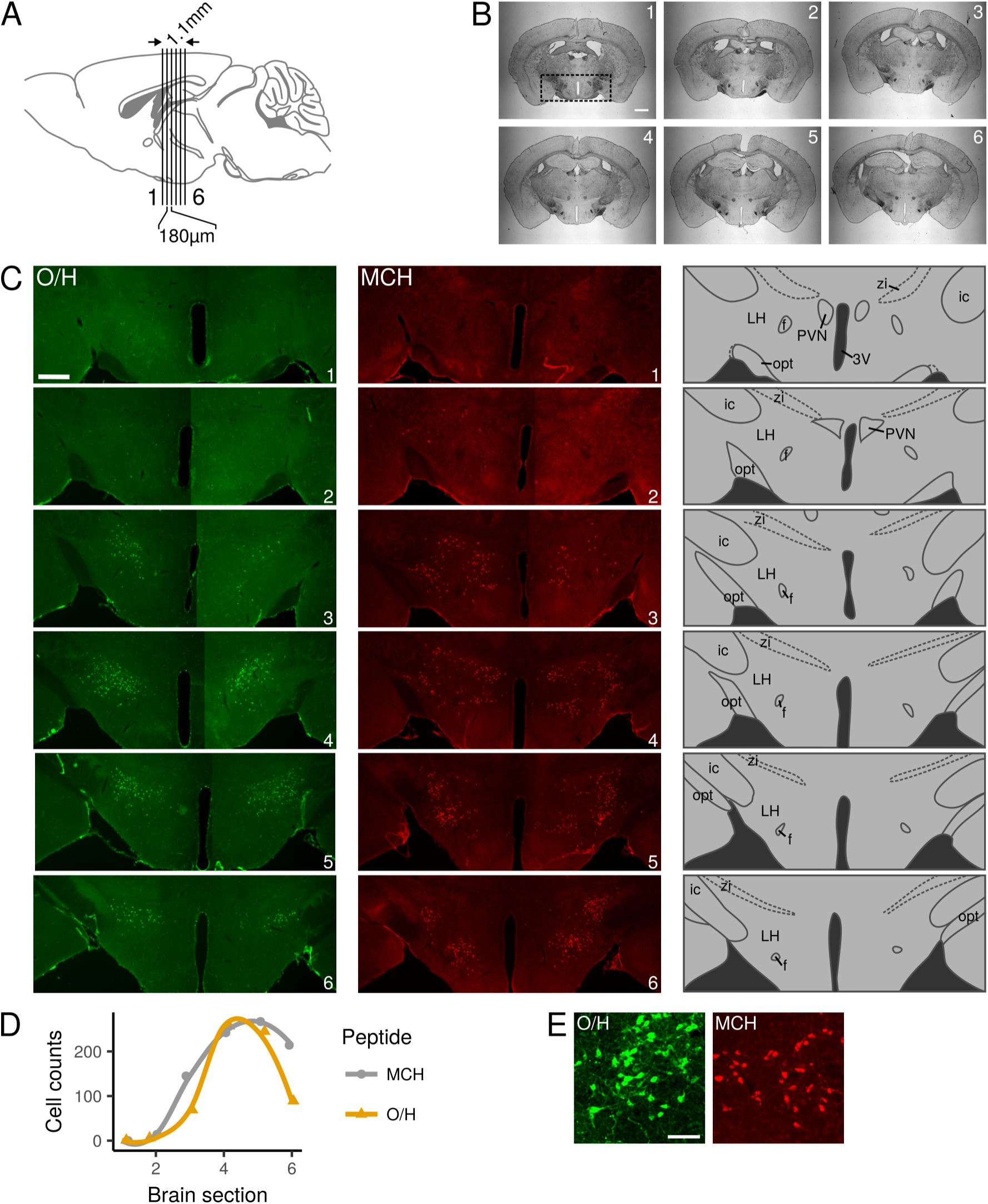
Cell counts in the lateral hypothalamus. (A) Each brain was sliced rostro-caudally into 30 μm-thick coronal sections. (B) The first section collected for immunofluorescence was the one at the caudal end of the optic chiasm, at the point where the optic tract begins its dorso-lateral trajectory into the brain (B1; ~0.94 mm caudal from bregma in Paxinos & Franklin, 2001). Then, every 6^th^ section (and its adjacent section) was collected in rostro-caudal sequence for a total of 2 sets of 6 sections per brain; thus, each set of brain sections were 180 μm apart, and the region studied spanned ~1.1 mm, which covers the area of orexin/hypocretin (O/H) expression in the lateral hypothalamus. The area delimited in (B1) is the same as that in (C1, O/H). (C) Brain sections in (B) were processed for immunofluorescence to reveal O/H-like immunoreactivity (left column), and the sections immediately adjacent to each of these were processed for melanin-concentrating hormone (MCH)-like immunoreactivity (middle column). Pictures from both sides of the brain were obtained; here, each image consists of two micrographs per brain section stitched together to show the full extent of the area studied at each coronal plane. The diagrams (right column) highlight the main anatomical landmarks (3V, third ventricle; f, fornix; ic, internal capsule; LH, lateral hypothalamus; opt, optic tract; PVN, paraventricular nucleus; zi, zona incerta). (D) Immunoreactive cells in each brain section were counted and plotted, as shown here for one animal. Notice the correspondence between section number (*x* axis) and brain sections as depicted in (A), (B), and (C). (E) Digital zoom of a group of cells in (C4). Individual immuno-labelled neurones can clearly be identified. Scale bar is 1 mm in (B), 500 μm in (C), and 100 μm in (E). The sagittal brain drawing in (A) was adapted from Paxinos & Franklin (2001).

For immunofluorescence, free-floating brain sections were rinsed in PBS, permeated with Triton (0.1% in PBS) for 15 min, incubated in glycine (1M in PBS) for 30 min, rinsed in PBS, blocked with albumin from bovine serum (BSA, 1% in PBS) for 30 min, and incubated overnight at 4°C with the primary antibody (see below) diluted 1:2000 in 1% BSA. Then, the sections were rinsed in PBS and incubated for 2 h at room temperature with the secondary antibody (1:400 in 1% BSA), rinsed in PBS, incubated in Hoechst (1:500 in PBS), mounted with FluorSave, and stored at 4°C.

The antibodies used were (i) rabbit anti-MCH antibody (raised against human, rat, and mouse MCH; Phoenix Pharmaceuticals H-070–47, lot 01329–1, Antibody Registry[RRID]:AB_10013632) followed by goat anti-rabbit Alexa Fluor 568 (Invitrogen A11011, RRID:AB_143157), or (ii) goat anti-O/H (raised against a peptide mapping at the C-19 end of human orexin-A; Santa Cruz Biotechnology sc-8070, lot I0611, RRID:AB_653610) followed by donkey anti-goat Alexa Fluor 488 (Invitrogen A11055, RRID:AB_2534102). In our experiments the expression pattern of MCH- and O/H-ir cells (Fig. 1C) was as reported previously (e.g. Broberger *et al.*, 1998; Hahn, 2010). Also, in preliminary experiments we performed double immunofluorescence for both MCH and O/H and confirmed that there was no overlapping of these peptides, which was as expected because orexin and MCH are two different cell populations (Broberger *et al.*, 1998). The findings reported below were the result of single immunofluorescence for O/H or MCH in adjacent brain sections.

### 2.3 Data collection and analysis

Images were obtained with an epifluorescence microscope (Leica DM4000B; Leica Microsystems, Wetzlar, Germany) fitted with a 5×/0.15 objective for counting ir cells (as in Fig. 1C) or a 1.25×/0.04 objective for full-section images (as in Fig. 1B). The latter were used to confirm that all sections were anatomically equivalent across brains at each of the six rostrocaudal levels per brain studied. Recall that the resolving power *R* of a lens is *R* = *λ*/2NA, where *λ* is the wavelength of light and NA is the numerical aperture. Thus, with e.g. red light (*λ* ≈ 600 nm) the 5× objective (NA = 0.15) that was used for counting cells can resolve up to 2 μm, which is more than enough to allow for the identification of individual O/H- or MCH-ir neurones (Fig. 1E). For counting, two images per brain section were captured, one for each side of the brain (Fig. 1C); thus, the number of immunoreactive cells per section reported here represent the total found in the full coronal plane, and it was unnecessary to align the brains along the sagittal plane because we did not analyse the medio-lateral distribution of these cells. We used ImageJ (Schneider *et al.*, 2012) with the plugin PointPicker (http://bigwww.epfl.ch/thevenaz/pointpicker/) to identify and count cells, and during this process the investigator was blind to the experimental condition of the images.

Statistical analyses were performed using R version 3.3.2 (R Core Team, 2016). Body weight in wildtype animals (Fig. 2A) was analysed independently of that in *db/db* mice (Fig. 2B). In each case a two-factor ANOVA followed by Tukey’s Honest Significant Difference method was used to test for differences in weight. Cell counts in Figs. 3B,C and 4B,C were analysed with two-way ANOVA tests, whereas one-way ANOVA tests were used for data in Fig. 5B,D,E. Data in Fig. 5F were analysed with ANCOVA. In all cases MCH data were tested independently of O/H data, and Tukey’s method was used for post-hoc multiple comparisons of means.

**Figure 2.**
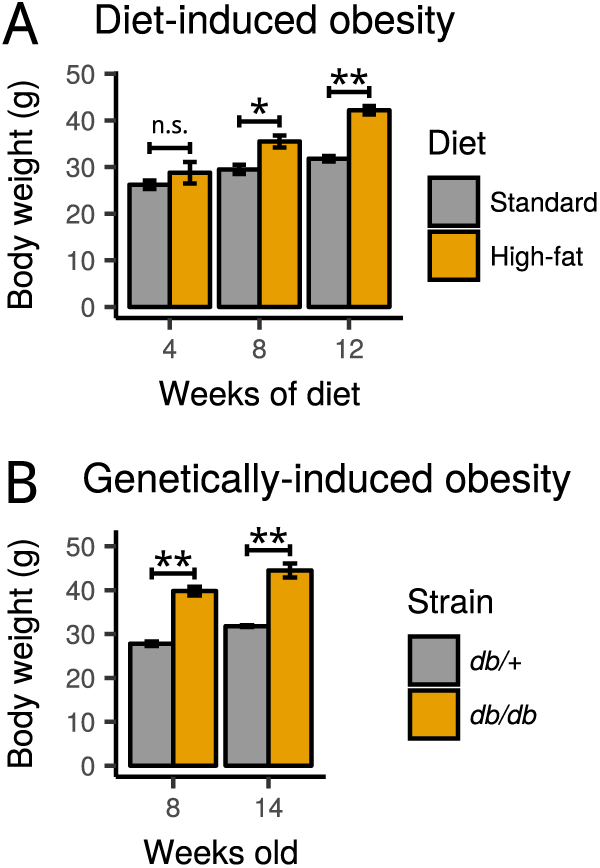
Body weight in animal models of obesity. (A) Wild-type mice were fed a diet high in fat starting at 4 weeks of age. Four weeks later, at age 8 weeks, their body weight was not significantly different from that seen in animals eating a standard diet. However, the animals that received the diet high in fat for 8 or 12 weeks (i.e. until 12 or 16 weeks of age) gained significantly more weight than control animals. (B) Genetically-obese *db/db* mice were significantly heavier than lean heterozygous (*db/+*) mice. n.s., not significant; \**p* = 0.039; \*\**p* ≪ 0.001. Each bar represents mean ± semof *n* = 4 animals.

**Figure 3.**
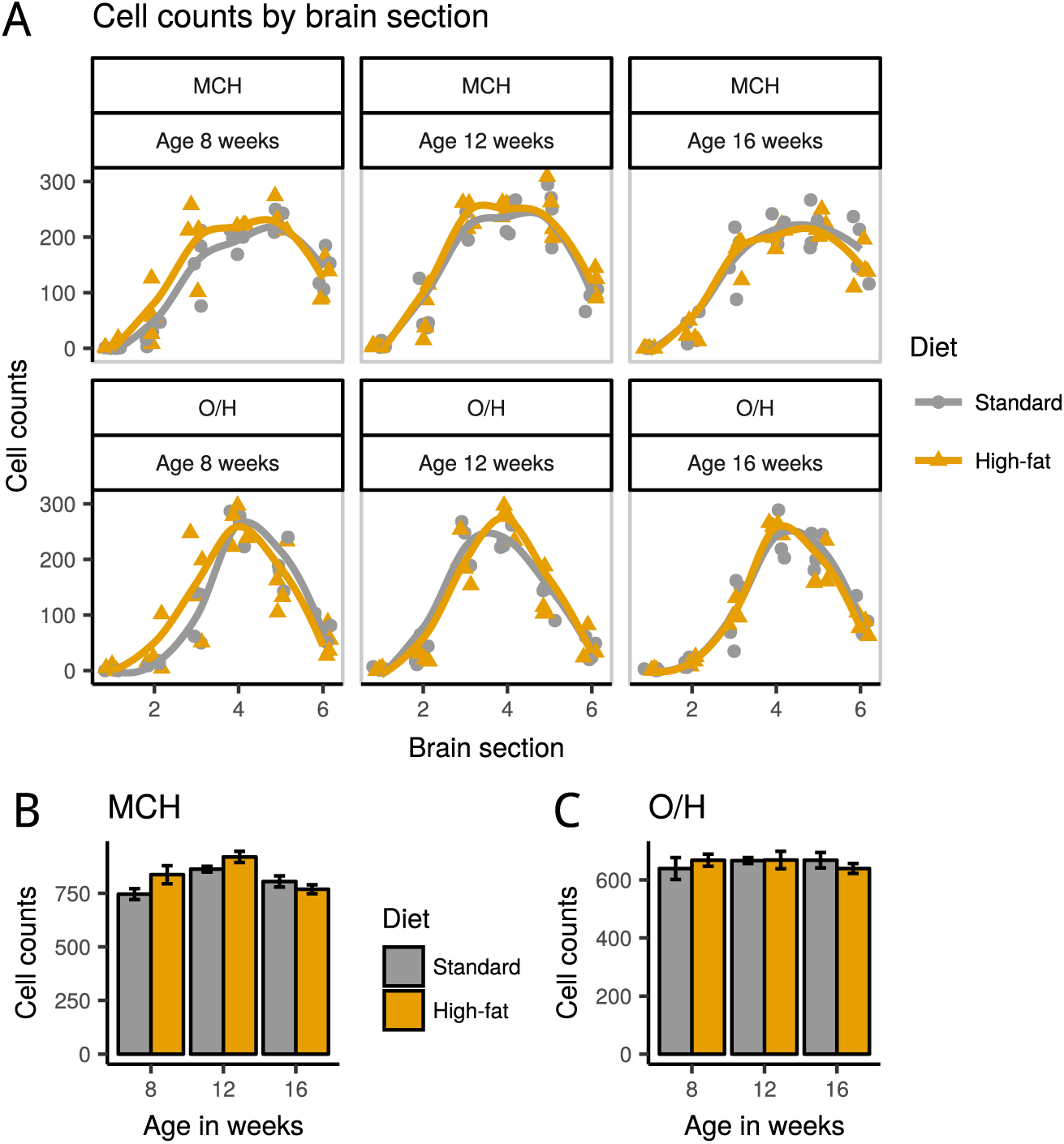
Melanin-concentrating hormone (MCH)- and orexin/hypocretin (O/H)-immunoreactive cell counts in diet-induced obese mice. (A) Cell counts by animal and brain section in mice fed a standard diet or a high-fat diet for 4, 8, or 12 weeks starting at 4 weeks of age. The total number of cells counted per animal is shown in (B) and (C), where each bar represents mean ± semcounts of *n* = 4 animals.

**Figure 4.**
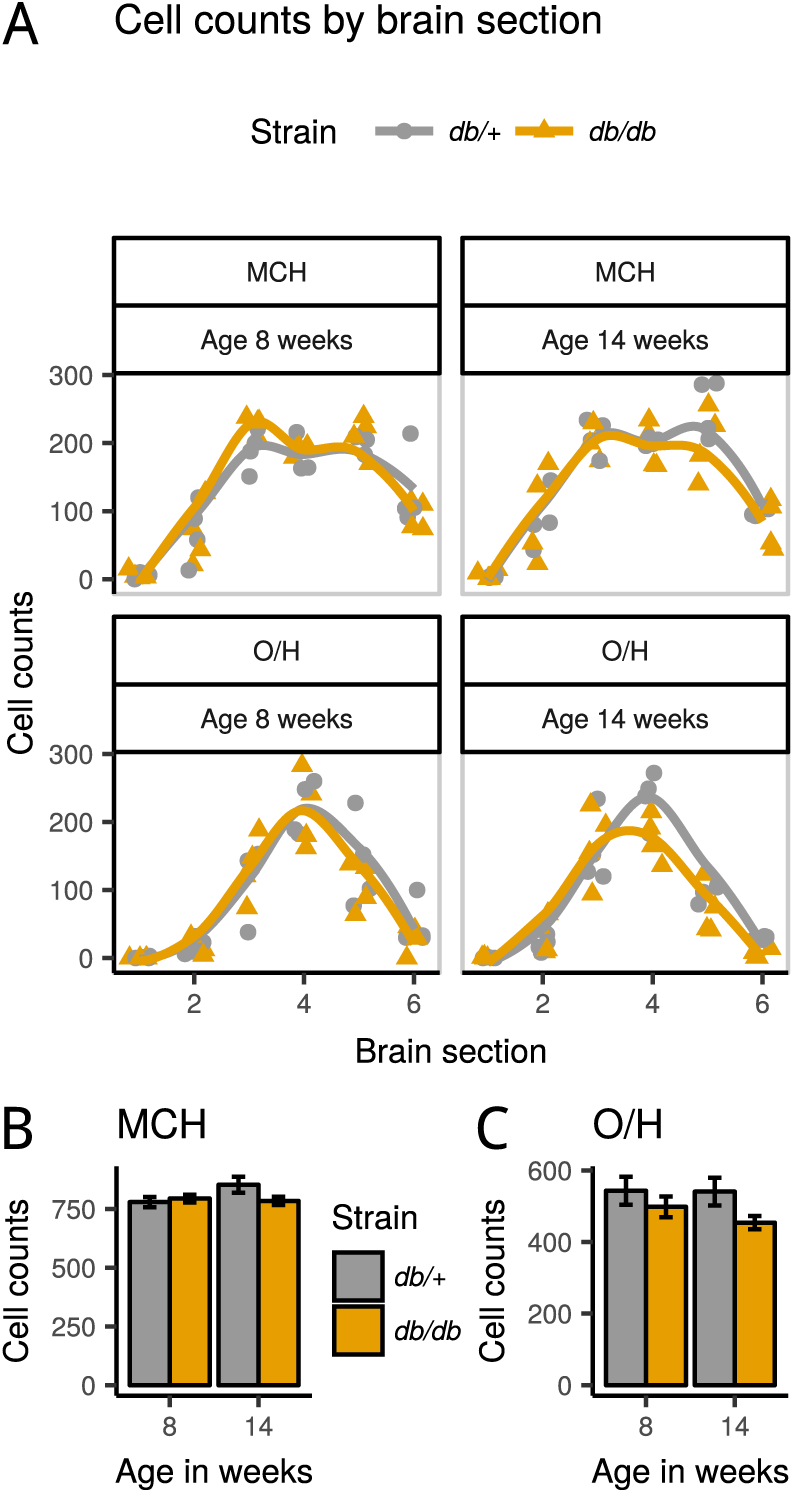
Melanin-concentrating hormone (MCH)- and orexin/hypocretin (O/H)-immunoreactive cell counts in genetically-induced obese *db/db* mice and lean *db/+* controls. (A) Number of immunoreactive cells by peptide, brain section, and animal. These data are presented in (B) and (C) as total MCH- and O/H-immunoreactive cells counted per animal. Each bar represents mean ± sem of *n* = 4 animals.

**Figure 5.**
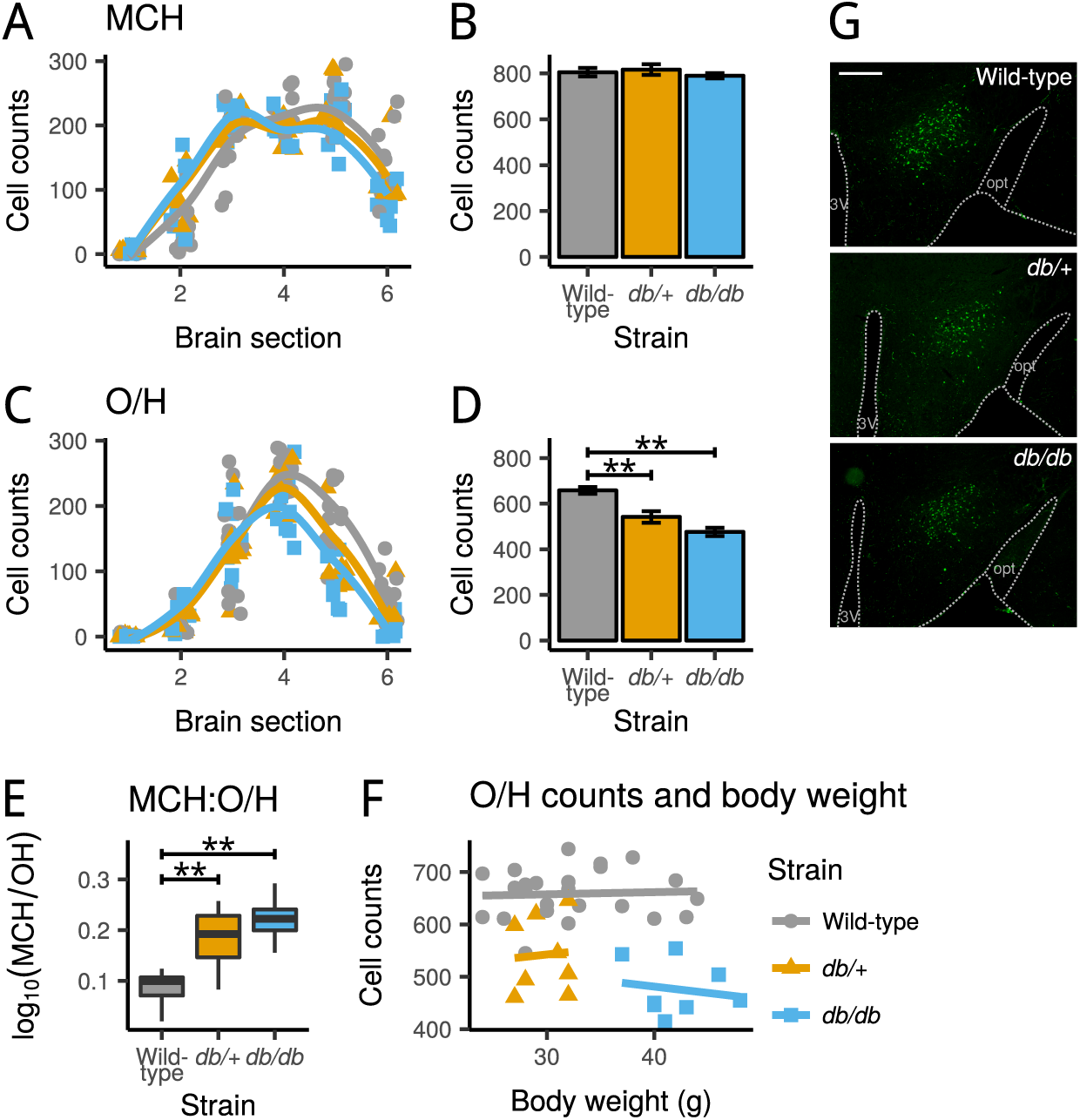
Cell counts in wild-type, *db/+*, and *db/db* mice (all fed on a standard diet). (A) Melanin-concentrating hormone (MCH)-ir cell counts by brain section, animal, and strain, and (B) total MCH cell counts by strain. (C) Orexin/hypocretin (O/H)- immunoreactive cell counts by brain section, animal, and strain, and (D) total O/H cell counts by strain. Bars represent mean±semimmunoreactive cell counts of *n* = 12animals (wild-type) or *n* = 8animals each (*db/+* and *db/db*). \*\**p* ≪ 0.001. (E) Ratio of MCH to O/H cells by strain. This ratio is expressed as the log10 of MCH counts divided by O/H counts. \*\**p* ≪ 0.001. (F) Relationship between O/H cell counts and body weight, by strain. The straight lines are linear fits. (G) Examples of immunofluorescence-labelled O/H cells in the lateral hypothalamus of wildtype, *db/+*, and *db/db* mice. A few landmarks are delineated for clarity (refer to Fig. 1). 3V, third ventricle; opt, optic tract. Scale bar, 500 μm.

## 3 Results

### 3.1 Animal models of obesity

To explore the possible effects of diet-induced obesity on MCH- and O/H-ir cells of the hypothalamus we fed a diet high in fat to wild-type mice for 4, 8, or 12 weeks, starting at 4 weeks of age. High-fat diet induced an increase in body weight that was significantly greater than that seen in animals fed on standard chow, and this effect depended on the duration (4, 8 or 12 weeks) of the exposure to this diet (interaction diet:duration, *F*(2,18) = 4.866, *p* = 0.021, two factor ANOVA).

Multiple comparisons of means (Fig. 2A) showed that it took HFD animals longer than 4 weeks to gain more weight than their standard-diet counterparts: four weeks after the diet protocols were introduced, mean (sem) body weight was 26.2 (0.9) g in the standard-diet group and 28.8 (2.3) g in the highfat diet group, a difference that was not significant (*n* = 4per group, *p* = 0.741, Tukey’s Honest Significant Difference). But animals that fed on HFD for 8 weeks became significantly heavier, and the weight gain was even more marked in animals that received that diet for 12 weeks. Mean (sem) body weight in the 8-week group was 29.5 (1.0) g in control (standard diet) and 35.5 (1.3) g in HFD animals (*n* = 4 per group, *p* = 0.039), and in the 12-week group it was 31.8 (0.6) g and 42.2 (0.9) g for standard and high-fat diet groups respectively (*n* = 4per group, *p* = 0.0002).

We also studied genetically-obese *db/db* mice to evaluate the effects of obesity on MCH- and O/Hir cell numbers independently of diet. Overall, *db/db* mice were much heavier than control *db/+* mice (strain, *F*(1,13) = 173.82, *p* = 6.7 × 10^−9^; age, *F*(1,13) = 21.73, *p* = 0.0004; two factor ANOVA), and there was no significant interaction between animal strain (*db/+* or *db/db*) and age (8 or 14 weeks old) (*F*(1,12) = 0.149, *p* = 0.706) which suggests that the marked difference in body weight was already established by the time the animals were 8 weeks of age (Fig. 2B). Mean (sem) body weight in 8-weeksold mice was 27.8 (0.5) g and 39.8 (1.0) g (*db/+* and *db/db* mice respectively, *n* = 4per group, *p* ≪ 0.001, Tukey’s Honest Significant Difference), and in 14- weeks-old mice it was 31.8 (0.2) g and 44.5 (1.6) g (*db/+* and *db/db, n* = 4per group, *p*≪0.001).

### 3.2 MCH- and O/H-ir cell counts in diet-induced obese mice

We counted MCH- and O/H-ir cells in the hypothalamus of wild-type animals that received standard diet or a diet high in fat for 4, 8, or 12 weeks starting at 4 weeks of age. Cell counts were from 6 coronal sections (numbered 1 to 6 in rostro-caudal sequence; see Fig. 1) per neuropeptide per brain, and adjacent brain sections were used for MCH or O/H immunostaining. The resulting data are shown in Fig. 3A. We hypothesized that the number of MCH- or O/H-ir cells was a function of high-fat diet and of the length of time the animals were exposed to the diet. Because the diet was introduced at 4 weeks of age in all experimental animals, the time these animals were exposed to high-fat diet was considered to be represented by their age.

We found that diet did not affect the number of MCH-ir cells, but there was a significant and unexpected difference in the number of MCH cells with age (diet, *F*(1,20) = 2.34, *p* = 0.142; age, *F*(2,20) = 7.94, *p* = 0.003; two-way ANOVA). Multiple comparisons of means revealed that this difference was on account of a greater number of MCH cells in 12- weeks-old animals (mean [sem] MCH cells was 791.5 [28.5] in 8-weeks-old mice, 891.4 [17.3] in 12-weeks-old mice, and 787.1 [16.9] in 16-weeks-old mice, *n* = 8 animals per group; Tukey’s Honest Significant Difference, 8 vs 12 weeks, *p* = 0.008; 12 vs 16 weeks, *p* = 0.006, 8 vs 16 weeks, *p* = 0.988; Fig. 3B).

Contrary to our expectations, the number of O/Hir cells did not change with diet or age (diet, *F*(1,20) = 0.001, *p* = 0.977; age, *F*(2,20) = 0.213, *p* = 0.810; two-way ANOVA; Fig. 3C). Thus, from a statistical point of view all wild-type animals studied had an equivalent number of O/H-ir neurones in the brain region analysed (mean [sem] O/H cells, 658.5 [9.6], *n* = 24 mice). In sum, high-fat diet had no effect on the number of MCH or O/H-ir cells. A transient increase in the number of MCH (but not O/H) neurones was observed at 12 weeks of age irrespective of the type of diet.

### 3.3 MCH- and O/H-ir cell counts in *db/db* mice

To investigate if obesity affected MCH and/or O/H cells independently of diet we counted MCH- and O/H-ir neurones in obese *db/db* mice and in lean *db/+* control mice 8 or 14 weeks old (Fig. 4A) following the same procedure described above (Fig. 1).

We did not observe any differences in MCH-ir cell numbers in these mice at the ages studied (strain, *F*(1,13) = 1.107, *p* = 0.312; age, *F*(1,13) = 1.587, *p* = 0.230, two-way ANOVA; Fig. 4B). Likewise, there were no significant differences in O/H-ir cell counts (strain, *F*(1,13) = 4.322, *p* = 0.058; age, *F*(1,13) = 0.544, *p* = 0.474; two-way ANOVA; Fig. 4C), although obese *db/db* mice seemed to have marginally fewer O/H cells than lean *db/+* controls (mean [sem] counts: *db/+* mice, 542.0 [25.4]; *db/db* mice, 476.1 [18.1]; *n* = 8 in each group).

### 3.4 O/H-ir cell counts in *db/db* vs wild-type mice

While diet did not affect the number of MCH or O/H cells, we observed that on average *db/db* animals seemed to have fewer O/H neurones than wild-type mice. Thus, we compared the number of O/H-ir cells between wild-type, *db/+*, and *db/db* mice to see if the *db* genotype alone could explain differences in cell counts (Fig. 5). To avoid confounding factors we excluded here animals fed a high-fat diet, so that only data from animals fed a standard diet were analysed (wild-type mice, *n* = 12; *db/+* mice, *n* = 8; *db/db* mice, *n* = 8). Animal strain had a very significant effect on the, number of O/H-ir (*F*(2,25) = 25.42, *p* = 9.5 × 10^−7^, one-way ANOVA; Fig. 5D) but not MCH-ir neurones (*F*(2,25) = 0.434, *p* = 0.653, one-way ANOVA; Fig. 5B). Multiple comparisons of means revealed that both *db/+* and *db/db* mice had fewer O/H cells than wild-type mice (wild-type vs. *db/+, p* = 0.0005; wild-type vs. *db/db, p* = 9×10^−7^; *db/+* vs. *db/db, p* = 0.077; Tukey’s Honest Significant Difference). The same effect was confirmed after analysing the individual ratio of MCH to O/H cells (Fig. 5E), which had an overall range of 1.048 to 1.959, meaning that all animals had more MCH-ir than O/H-ir cells. But there was a very significant effect of strain on this ratio (*F*(2,25) = 21.63, *p* = 3.5×10^−6^, one-way ANOVA, calculated using the log10 of MCH counts divided by O/H counts), and this was because both *db/+* and *db/db* mice had a much greater ratio of MCH to O/H neurones than wild-type mice (wild-type vs. *db/+, p* = 0.0006; wild-type vs. *db/db, p* = 4.3×10^−6^; *db/+* vs. *db/db, p* = 0.198, Tukey multiple comparisons of means).

Finally, we investigated if the number of O/H cells was related to body weight using data from all animals (*n* = 40). Mice become overweight within days after ablating O/H cells (González *et al.*, 2016) which shows that the number of orexin cells directly affects body weight, but it is not known if such relationship holds true for less-extreme cases. Here, strain was necessarily included as a co-factor because of the strong link between body weight and strain (Fig. 2). As illustrated in Fig. 5F, there was a marked effect of strain on O/H cell numbers (which was already described above) but no evidence of a correlation between body weight and the number of O/H cells (strain, *F*(2,36) = 39.59, *p* = 8.1×10^−10^; body weight, *F*(1,36) = 0.014, *p* = 0.907; analysis of covariance). In fact only a very small percentage of the variation in body weight was accounted for by O/H cell counts. This percentage was 0.3% in wild-type animals (*R*^2^ = 0.0026), 0.5% in *db/+* mice (*R*^2^ = 0.0049), and 3% in *db/db* mice (*R*^2^ = 0.028).

## 4 Discussion

Diet has direct implications for body weight, yet the links between the two are still not fully understood. Recent studies have demonstrated that a diet high in fat triggers histopathological changes in the arcuate nucleus of the hypothalamus that include inflammation, gliosis, apoptosis, and synaptic reorganization (Moraes *et al.*, 2009; Thaler *et al.*, 2012; Buckman *et al.*, 2013; Horvath *et al.*, 2010). These changes seem to predominantly affect specific cell types directly involved in energy homoeostasis and are thought to be a leading cause of metabolic imbalance and obesity (see Jais & Brüning, 2017, for a recent review). Apoptosis was also observed in the lateral hypothalamus in response to high-fat diet (Moraes *et al.*, 2009), but we do not know which cell types are affected. Two cell populations in the lateral hypothalamus, orexin/hypocretin and melanin-concentrating hormone neurones, are essential for appetite and metabolism regulation (Burdakov *et al.*, 2013). Thus, we hypothesized that HFD would decrease the number of O/H and/or MCH cells. Contrary to our expectations we found no differences in O/H- or MCHir cell numbers in mice that received HFD for up to 12 weeks compared to mice that received standard chow (Fig. 3) despite the marked difference in body weight between these groups (Fig. 2). These results are in agreement with those by Lemus *et al.* (2015), who also found that high-fat diet did not affect the number of O/H cells in mice (they did not quantify MCH cells), and more generally with those by Namavar *et al.* (2012), who did not see a change in total neurones in the hypothalamus after high-fat diet. By contrast, we found that genetically-obese *db/db* mice had significantly fewer O/H-ir (but not MCH-ir) cells (Fig. 5). Lean heterozygous *db/+* animals also had fewer O/H-ir cells than wild-type mice but the effect was less marked.

We counted immuno-labelled cells in the lateral hypothalamus ensuring that brain sections were anatomically equivalent between subjects (Fig. 1). Whereas within-subject brain sections were far enough apart from each other to avoid double counting, at the same time our experimental design guaranteed that a comprehensive set of MCH- and O/Hir cell populations was sampled along the full rostrocaudal range. We also ensured that we followed a careful quantitative analysis of our data. We note that Yamamoto *et al.* (1999) reported no changes in O/H-ir cell numbers in *db/db* mice. However, they seemingly based their conclusions on a qualitative description of two micrographs (their Figure 4). If we had followed that same approach we would have reached similar conclusions to theirs because at first glance there were no striking differences in O/H-ir cells between strains in our immunofluorescence images (Fig. 5G). In that same paper Yamamoto *et al.* (1999) found a significant decrease in the expression of *prepro-O/H* mRNA in *db/db* mice compared to wildtype mice using a semi-quantitative approach. These latter findings are consistent with the ones we are reporting here.

The reduced number of O/H-ir cells that was seen in obese *db/db* mice may be due to a decrease in the expression of the peptide below detection levels and/or to a loss of peptide-expressing cells. Either way our findings most likely represent a reduction in available hypothalamic O/H in these mice. Genetically-obese *db/db* mice and diet-induced obese mice are both insensitive to leptin and have high plasma levels of the hormone. In the former this is because of a mutation in the long form of the leptin receptor that makes cells insensitive to leptin (Lee *et al.*, 1996), whereas in the latter leptin resistance is presumably the result of high-fat diet-induced hypothalamic injury (Münzberg *et al.*, 2004). If a failure in leptin signalling was responsible for the changes in O/H in *db/db* mice, why were diet-induced obese mice not affected? Perhaps O/H cells are sensitive to defects in leptin signalling mostly during development. High-fat diet in our experiments was introduced at 4 weeks of age, well ahead of the time at which O/H and MCH cells develop, which is between embryonic days 9–14 (Croizier *et al.*, 2010), whereas *db/db* mice are already affected by the leptin receptor mutation at those ages. It is interesting to note that both O/H and MCH mRNA expression increases in pups of rats fed a high-fat diet during pregnancy (Beck *et al.*, 2006; Chang *et al.*, 2008), suggesting that these peptidergic systems are sensitive to changes in diet before birth.

Differences between diet-induced and genetically-induced obese mice may also be explained by the extent of the leptin resistance because it has been suggested that leptin resistance in animals that are fed a diet high in fat develops only in the arcuate nucleus and not in other hypothalamic areas (Münzberg *et al.*, 2004), whereas leptin-signalling defects in *db/db* mice are not region specific. It is also possible that the observed differences in O/H-ir between animal models were due to a more general effect on brain development because the brains of leptin-resistant *db/db* mice and of leptin-deficient *ob/ob* mice are significantly smaller than expected (Vannucci *et al.*, 1997; Steppan & Swick, 1999). Leptin replacement in the latter seems to directly increase the number of brain cells (Steppan & Swick, 1999). However, two observations argue against this and suggest instead a specific effect of the *db* genotype on O/H neurones: (i) a non-specific general developmental effect would presumably affect both MCH and O/H neuronal populations, yet only O/H-ir cells were affected in *db/db* mice (Fig. 5); and (ii) by contrast to obese *db/db* mice, lean *db/+* mice are not different physiologically or anatomically from wild-type mice (Hummel *et al.*, 1966), yet in our studies *db/+* mice also had fewer O/H-ir neurones than wild-type mice (Fig. 5).

### 4.1 Conclusions

We found fewer hypothalamic O/H-ir cells in *db/db* and *db/+* mice than in wild-type mice, and this effect was only explained by the *db* genotype and not by high-fat diet intake, age, or body weight. The number of MCH-ir cells was not affected by genotype or diet, but we saw a moderate and transient increase in these at 12 weeks of age. Our findings show that lateral hypothalamic neurones that are crucial for the mainte-nance of energy homoeostasis are affected differently in diet-induced obese animals and in mice that are genetically obese, and support the notion that highfat diet intake does not affect equally all components of the network that keeps metabolism in tune.

## Author contributions

All authors had full access to all the data in the study. Study concept and design: J.A.G. and J.H.M.P. Acquisition, analysis, and interpretation of data: J.A.G. Drafting of the manuscript: J.A.G. Critical revision of the manuscript: J.A.G. and J.H.M.P. Obtained funding: J.A.G. and J.H.M.P.

## Acknowledgements

Work funded by a Marie Curie Intra-European Fellowship (FP7-PEOPLE-2009-IEF Project number 255559 to J.A.G.). The authors declare no conflicts of interest.

## Abbreviations

HFD: high-fat diet
ir: immunoreactive
MCH: melanin-concentrating hormone
O/H: orexin/hypocretin

